# EpiReasoner: An Integrated Artificial Intelligence Framework for Phenotype-to-Genotype Reasoning in Plant Epidermal Development

**DOI:** 10.64898/2026.05.13.724792

**Authors:** Haoyang Zhang, Lunzeng Huang, Kun Ning, Jing He, Yong He, Jingquan Yu, Zhong-Hua Chen, Zhangjian Hu, Xuping Feng

## Abstract

Achieving high-throughput and precise phenotypic quantification and imaging modalities of stomatal and epidermal cells across diverse species remains a primary bottleneck in elucidating the mechanisms of stomatal dynamics, epidermal patterning, and environmental adaptation of plants. Here, we developed EpiReasoner, an artificial intelligence framework comprising a vision module, EpiVision, and a knowledge-based reasoning module, EpiBrain, for the quantitative phenotypic analysis and domain-specific knowledge reasoning of stomatal complexes and pavement cells in plants. Operating across bright-field, scanning electron microscopy, and differential interference contrast modalities, EpiVision achieves precise instance segmentation in various monocotyledonous, dicotyledonous, and fern species. Its performance significantly surpasses current state-of-the-art models. Moreover, we defined 23 quantitative indices describing stomatal cell morphology and spatial distribution. For domain-specific tasks such as phenotype prediction, genotype deduction, and molecular mechanism reasoning, EpiBrain demonstrates a human preference rate significantly higher than that of general-purpose large language models, including GPT-5 and Claude Sonnet 4. The application of EpiReasoner to phenotypic data of stomatal density derived from a tomato natural population of 170 accessions successfully identified a major quantitative trait locus on chromosome 8. The candidate gene, *SKP1-interaction partner 19L* (*SKIP19L*), encoding an F-box family protein, exhibited severe allele frequency drift during tomato domestication, which is highly consistent with the adaptive trend of reduced stomatal density under artificial selection. EpiReasoner provides a novel paradigm that unifies visual phenomics and knowledge-driven reasoning for the biology of stomata and pavement cells, thereby significantly accelerating scientific discovery in plant science.

## Main

The leaf epidermis serves as a critical physical and physiological barrier, essential for plant survival and environmental adaptation. The leaf epidermis predominantly include pavement cells, guard cells, subsidiary cells, and trichomes. Among these, pavement cells occupy the majority (typically > 80-95%) of the epidermal area, functioning primarily to provide mechanical support, maintain tissue integrity, and prevent excessive water loss and external pathogen invasion^1,2^. Stomatal complexes, formed by two guard cells and sometimes two subsidiary cells (in the case of grasses and other monocots), serve as vital gates for precisely regulating the gas exchange required for plant transpiration and photosynthesis^3,4^.

Given that stomatal complexes exhibit significant morphological and spatial distribution variations across different plant taxa, the precise quantification of their number, arrangement, status of opening/closure, and density/index is a prerequisite for elucidating the mechanisms of epidermal development, stomatal regulation, and environmental adaptation of plants. To this end, machine learning technologies have recently demonstrated immense potential in enhancing the throughput of plant phenotyping and significant progress has been achieved in the recognition and analysis of stomatal-related phenotypes^5^. Examples include high-throughput stomatal identification and counting based on convolutional neural networks^6^, the precise extraction of stomatal morphology and stomatal pore area based on instance segmentation^7,8^, guard cell homeostasis models^9,10^, and the investigation of circadian rhythms in in-situ stomatal movements based on video tracking^11^. Although higher-order methods, such as computational geometry and network graph theory, have been introduced to quantify their topological features^12,13^, such analyses remain highly dependent on the segmentation efficacy of the target objects. Overall, these studies predominantly focus on methodological design and optimization for single imaging modalities or individual species, restricting the generalizability of the models across different imaging conditions and diverse groups of plant species.

With the breakthroughs in universal visual models and architectures, cell segmentation methodologies are evolving from task-specific approaches toward universal solutions^14,15^. Mainstream universal cell segmentation models, primarily driven by medical or animal research data, have been successfully adapted for biomedical applications through domain-specific fine-tuning on annotated target datasets^16–18^. When directly applied to the unique low contrast and extreme cellular geometric variations characteristic of plant epidermal microscopic materials, these universal models frequently encounter substantial domain shift issues. Therefore, the design of targeted fine-tuning or domain adaptation strategies to manage the complex scenarios of cell types of plant epidermis, such as stomata, necessitates further exploration.

The precise acquisition of stomatal phenotypic data is a prerequisite for formulating scientific questions and research hypotheses. Throughout this process, the comprehensive synthesis of the extensive literature is pervasive, providing foundational support for experimental design, data interpretation, and mechanistic elucidation. However, confronted with the exponentially accumulating publications in the plant sciences, researchers are experiencing severe information overload, making it challenging to rely on traditional approaches for effective information retrieval, core information summarizing, and in-depth reasoning. Recently, large language models (LLMs) have exhibited major breakthroughs in natural language processing, knowledge retrieval, and complex logical reasoning, providing a novel paradigm for mining scientific mechanisms based on massive corpora and achieving broad application across multiple vertical domains^19,20^. The plant science domain has also recently initiated active exploration into the application of LLMs. For instance, PlantScience.ai integrates knowledge graphs and retrieval-augmented generation technologies to construct a comprehensive intelligent question-answering assistant for plant science^21^, while PlantGPT fine-tunes open-source models using large-scale literature and phenotype-gene annotation data to focus on plant functional genomics reasoning^22^. Nevertheless, these models are primarily oriented toward macroscopic, general plant science contexts; the knowledge granularity and reasoning depth of their built-in knowledge bases regarding specific domains (e.g. stomatal function, phenotypic traits, stress responses, and epidermal development) remain insufficient to directly support the in-depth scientific reasoning required for stomata specific questions.

Therefore, this study proposes EpiReasoner, an artificial intelligence (AI) system integrating visual phenotypic extraction and knowledge reasoning to address stomata-related questions in plants. The main objectives were: 1) to establish domain-specific fine-tuning strategies tailored for stomata and epidermal cells, thereby enabling the robust cross-species and cross-imaging modality generalization of large models in acquiring stomata phenotypes from diverse biological materials while maintaining universal capabilities; 2) to provide systematized intelligent assistance for experimental design, data interpretation, and mechanistic reasoning in conjunction with LLMs, thereby enhancing research efficiency and depth; 3) to systematically validate the comprehensive performance of EpiReasoner using a natural population of tomato (*Solanum lycopersicum*).

## Results

### EpiReasoner: A phenotypic profiling system from multimodal segmentation to knowledge reasoning

An overview of our AI system EpiReasoner is illustrated in Fig. 1. The system comprises a vision module EpiVision and a knowledge reasoning module EpiBrain (Fig. 1a). Specifically, the EpiVision module employs the vision foundation model Segment Anything Model 3 (SAM3)^23^ as its architecture. Following pre-training data of pavement cells and stomatal complexs derived from a dicotyledonous species tomato (*Solanum lycopersicum*) and a monocotyledonous species wheat (*Triticum aestivum*), it achieves the prompt-based segmentation and recognition of stomatal complexes and pavement cells across 3 imaging modalities and 3 distinct sample preparation conditions of the two plant species. Based on the generated high-precision segmentation masks, this module extracts quantitative indices describing the geometric and morphological parameters and boundary topological complexities of stomata and pavement cells. Furthermore, the spatial distribution patterns of stomata and pavement cells are quantified through the construction of cellular spatial adjacency graphs (Fig. 1b). The EpiBrain knowledge reasoning module is a specialized LLM constructed specifically for the domain of plant epidermal cell biology. This module utilizes a parameter-frozen Qwen3-32B model^24^ as its foundation, which is efficiently fine-tuned using Low-Rank Adaptation (LoRA)^25^ technology on a high-quality domain question-answering dataset comprising 173,335 entries compiled from literature. This configuration enables the module to maintain general language comprehension capabilities while possessing the capacity for in-depth knowledge retrieval and logical reasoning. The key areas are the developmental processes, environmental adaptations, cellular functions, and genetic regulatory mechanisms of plant epidermal cell types, with an emphasis on stomatal complex.

**Fig. 1.**
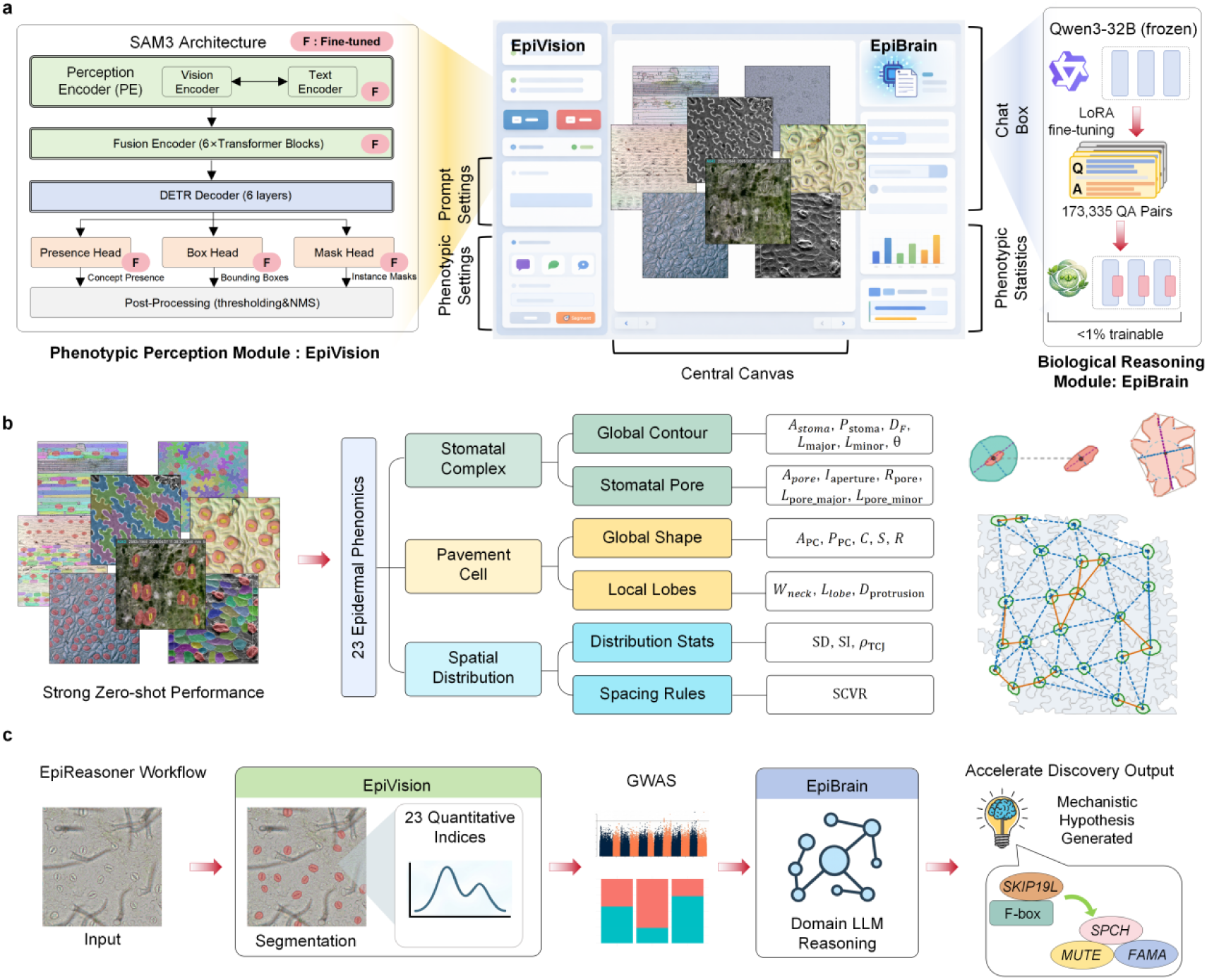
Architecture and overview of the EpiReasoner framework and phenotypic quantification scheme. **a**, EpiReasoner framework integrates EpiVision for visual feature extraction with EpiBrain for knowledge-driven reasoning of plant epidermal traits. **b**, EpiVision exhibits strong zero-shot segmentation performance across multimodal imaging data, enabling the quantitative extraction of 23 phenotypic indices. These include morphological metrics for stomatal complexes and pavement cells, as well as their spatial distribution topologies. **c**, The EpiReasoner scientific discovery workflow. The integration of EpiVision-derived quantitative indices with GWAS results empowers EpiBrain to perform domain-specific LLM reasoning, thereby accelerating the generation of mechanistic hypotheses (e.g., the SKIP19L genetic regulatory network).

Under the EpiReasoner framework, high-throughput phenotypic traits extracted by EpiVision can be seamlessly integrated into genome-wide association studies (GWAS) as high-fidelity input phenotypes. The candidate quantitative trait loci (QTLs) identified by GWAS are then sent into EpiBrain for deep domain reasoning, facilitating the efficient generation of mechanistic hypotheses. Our new pipeline accelerates the closed-loop research trajectory from precise phenotypic quantification to functional gene mining, establishing a new paradigm for data-driven genetic discovery.

### Segmentation of stomatal complexes and pavement cells with EpiVision

Precise instance segmentation is a crucial prerequisite for achieving automated plant phenotyping. Initially, we constructed a dataset of leaf epidermis samples of tomato and wheat plants utilizing bright-field microscopy and completed instance-level annotation. This dataset encompasses 5,419 pavement cells, 1,471 stomatal complexes, and 629 stomatal pores from the tomato samples, alongside 1,780 pavement cells, 426 stomatal complexes, and 51 stomatal pores from the wheat samples (Fig. 2a). We trained EpiVision, based on the vision foundation model SAM3, to address segmentation problems and compared it with three other representative deep learning architectures Mask R-CNN^26^, OneFormer^27^, YOLO26-Seg^28^. Quantitative evaluation results based on 6 metrics indicate that the comprehensive segmentation performance of EpiVision surpasses that of the other three architectures (Fig. 2b). Notably, the Aggregated Jaccard Index (AJI) was improved by 51.28%, 32.96%, and 17.00% in EpiVision compared with Mask R-CNN, YOLO26-Seg, and OneFormer, respectively. Comparison of the segmentation efficacy of four models across different target categories (Fig. 2c) confirmed that EpiVision demonstrates superior segmentation capacity when processing dicotyledonous pavement cells with complex topological boundaries and tiny stomatal pores.

**Fig. 2.**
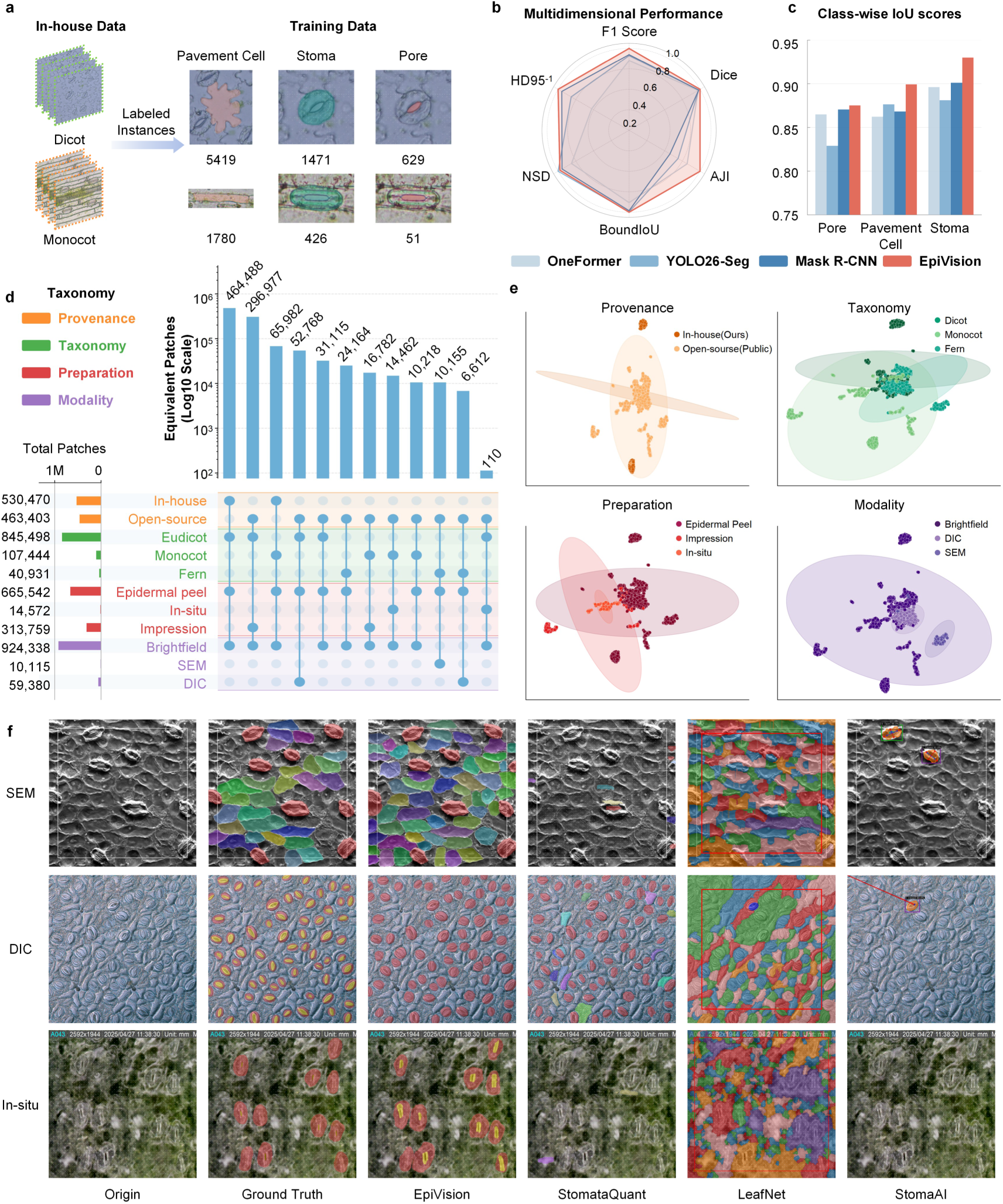
Performance evaluation of EpiVision. **a**, Composition of the training dataset. A total of 465 microscopy samples annotated for pavement cells, stomata, and pores were collected from 170 tomato accessions (dicotyledons) and 210 wheat lines (monocotyledons). **b**, Comparative segmentation performance of four models on in-house data. Performance was evaluated utilizing six core metrics. **c**, Segmentation results of the four models across distinct cell types. Performance was evaluated utilizing class-wise Intersection over Union. **d**, Overview of the evaluation dataset, EpiDataset, which comprises three microscopy acquisition modalities and three sample preparation protocols, spanning major plant taxa. **e**, Visualization of domain shifts within the feature space of EpiDataset. Dimensionality reduction based on Uniform Manifold Approximation and Projection reveals discrepancies across training and evaluation datasets, species, sample preparation methods, and imaging modalities. **f**, Segmentation efficacy of the four segmentation tools under scanning electron microscopy, differential interference contrast, and in-situ scenarios.

We then conducted significant phenotypic investigations of the leaf epidermis involving diverse imaging modalities and phenotypic variations across species. To comprehensively evaluate the generalization capability of EpiVision in authentic complex scenarios, this study integrated our proprietary dataset with 13 open-source databases^5,29^ (Supplementary Table 1) to construct the leaf epidermal dataset named as EpiDataset (Fig. 2c). EpiDataset comprises extensive samples of dicotyledonous, monocotyledonous, and fern plants; sample preparation techniques including epidermal peeling, impression methods, and *in-situ* imaging. The imaging modalities encompass bright-field, scanning electron microscopy (SEM), and differential interference contrast (DIC). Standardized cropping and stratified random sampling were performed on the raw data (see Methods). Subsequently, the original SAM3 was utilized to extract high-dimensional latent space features from the sampled patches, and Uniform Manifold Approximation and Projection (UMAP)^30^ was employed for feature dimensionality reduction. The UMAP scatter plot demonstrates that different origins, species, preparation methods, and imaging modalities exhibit significant clustering isolation within the deep feature space (Fig. 2d). This data distribution shift poses a challenge for evaluating the cross-domain generalization capability of vision segmentation models.

Furthermore, we compared EpiVision with three published computational models in the field of plant epidermal analysis (LeafNet, StomataQuant, and StomaAI). These three baseline tools exhibit differences in their design objectives and functional coverage scopes: StomaAI can identify stomatal complexes and stomatal pores, but lacks the capability to segment pavement cells; LeafNet can extract stomata and pavement cells without resolving stomatal pores; whereas StomataQuant supports the segmentation of all epidermal features, but its performance on *in-situ* and scanning electron microscopy samples was suboptimal. Visual qualitative analysis intuitively demonstrates that when confronting significant data distribution shifts, the performance of the respective models showed some variation (Fig. 2e). In cell segmentation tasks, the three models exhibit varying degrees of degradation in segmentation performance when processing samples under SEM and DIC images, or unseen species with complex epidermal features (such as ferns). In contrast, EpiVision consistently achieves comparatively precise instance segmentation of stomatal complexes, stomatal pores, and pavement cells across multiple heterogeneous imaging modalities.

### Construction and validation of quantitative indices for the morphology and spatial distribution of stomata and pavement cells

After segmentation with EpiVision, our system extracted a comprehensive set of 19 quantitative indices describing the morphology of stomata and pavement cells, and 4 quantitative indices characterizing spatial distribution (Fig. 1c and Fig. 3a, Supplementary Tabel 3). For the highly irregular pavement cells, we constructed the convex hull, determined the largest inscribed circle, and identified local curvature extrema along the cell boundary (specifically the positive and negative curvature peaks). These analyses enabled the extraction of 8 phenotypic features characterizing their morphological complexity.

**Fig. 3.**
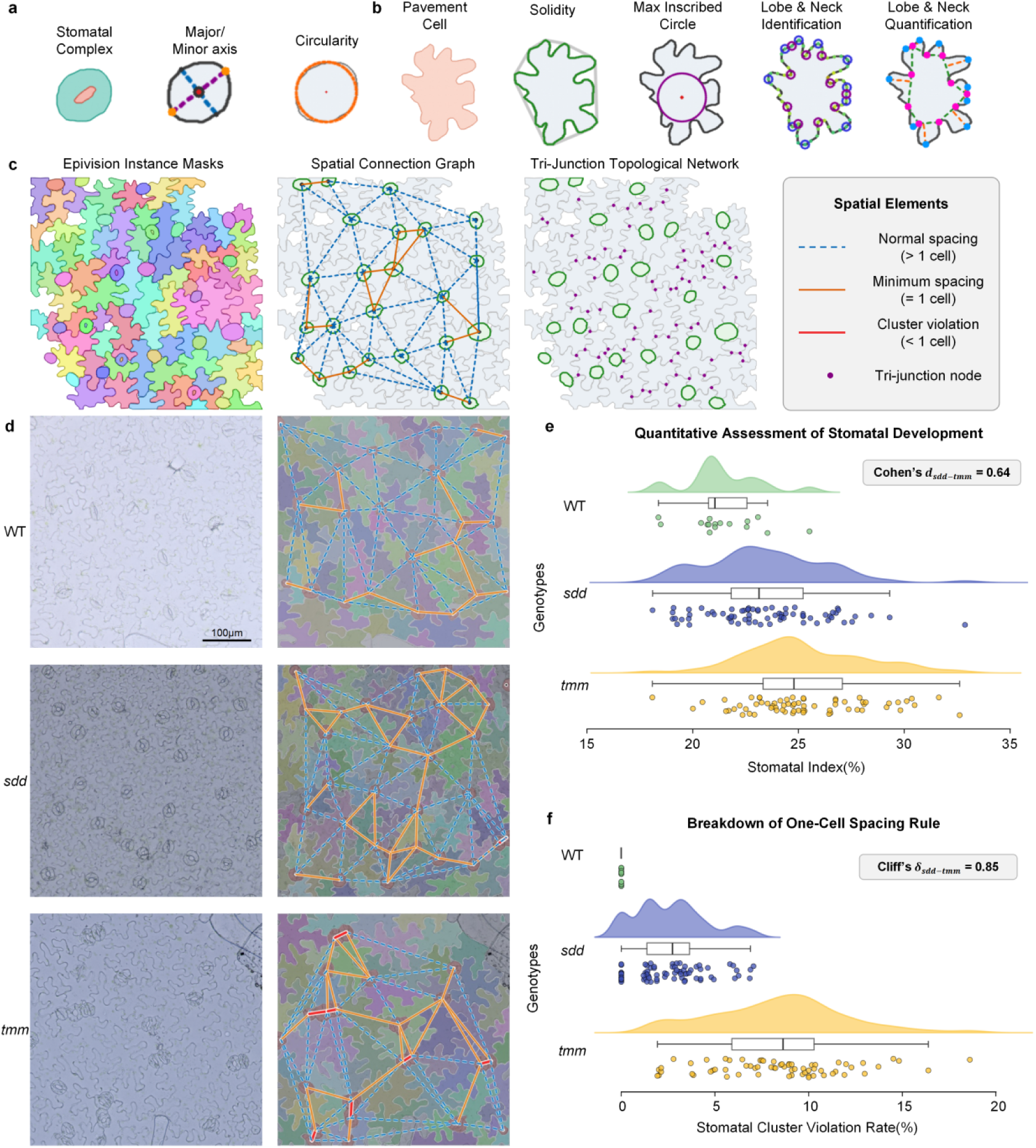
Quantitative metrics for stomatal and pavement cell phenotypes, and computational approaches for modeling epidermal organization and stomatal spatial distribution. **a**, Quantitative characterization methods for stomatal morphology. **b**, Quantitative characterization methods for pavement cell morphology. **c**, Delineation of the spatial distribution between stomata and pavement cells. A two-dimensional spatial connectivity network of stomata is constructed based on the one-cell spacing rule (middle; blue dashed lines and solid orange lines denote stomatal intervals greater than one and equal to one pavement cell, respectively, both indicating rule compliance), and a three-way junction network of pavement cells is extracted to characterize boundary assembly features (right). **d**, Spatial distribution models of stomata and pavement cells in wild-type and mutant (*sdd*, *tmm*) samples computed utilizing the Stomatal Cluster Violation Rate (SCVR). The red lines highlighted connections in the central topological graph capture and localize abnormal stomatal clusters with an interval of zero (indicating rule violation). **e**, Variances in the Stomatal Index (SI) among wild-type and mutant (*sdd* and *tmm*) samples. **f**, Variances in the SCVR among wild-type and mutant (*sdd* and *tmm*) samples.

In addition to the morphological analysis of individual cells, our framework further constructs a cellular spatial connectivity map to characterize the spatial organization patterns of stomata and pavement cells (Fig. 3c), introducing an index designated as the Stomatal Cluster Violation Rate (SCVR). The algorithm quantifies stomatal distribution by tracking the number of pavement cells intervening between adjacent stomata. To evaluate the capacity of SCVR at the biological resolution, we generated the homologous mutants *sdd* (*Stomatal Density and Distribution 1*) and *tmm* (*Too Many Mouths*) in tomato and applied this analytical pipeline to wild-type and the two mutant samples (Fig. 3d and e). *SDD1* encodes a subtilisin-like protease responsible for processing and generating extracellular signals that inhibit adjacent cells from entering the stomatal lineage^31,32^. *TMM* encodes a receptor-like protein that acts as a signal-specific receptor, forming a complex with ERECTA family kinases to recognize these signals, thereby restricting the excessive proliferation of stomatal precursor cells^33,34^. Pairwise Mann-Whitney *U* tests conducted across all samples revealed that although the Stomatal Index (SI) could reflect differentiation variances among genotypes (Kruskal-Wallis test, *P* < 0.001), its capacity to distinguish the spatial distribution patterns of stomata between *sdd* and *tmm* mutants was limited. In contrast, the SCVR proposed in this study was capable of significantly differentiating the phenotypic discrepancies between *sdd* and *tmm*. For the tomato wild-type samples, epidermal cells strictly adhered to the classic one-cell spacing rule^33^, yielding an SCVR of 0. The *sdd* mutant primarily exhibited significantly increased stomatal density, whereas the *tmm* mutant severely violated the one-cell spacing rule by forming numerous stomatal clusters. The comparison of SCVR between *sdd* and *tmm* mutants yielded a Cliff’s delta of 0.85, significantly outperforming the discriminative capacity of the SI (Fig. 3e and f). These results demonstrate that our proposed SCVR can precisely distinguish the extents of cellular quantity proliferation versus spatial arrangement alteration.

### EpiBrain for knowledge reasoning of the plant epidermis

To address the massive and fragmented data within the plant stomatal domain, we developed a specialized virtual scientist based on LLM reasoning, designated as EpiBrain (Fig. 4a). This system integrates domain-specific expertise spanning plant phenomics, molecular biology, developmental biology, plant physiology and stress biology, thereby assisting researchers in accelerating scientific discovery.

**Fig. 4.**
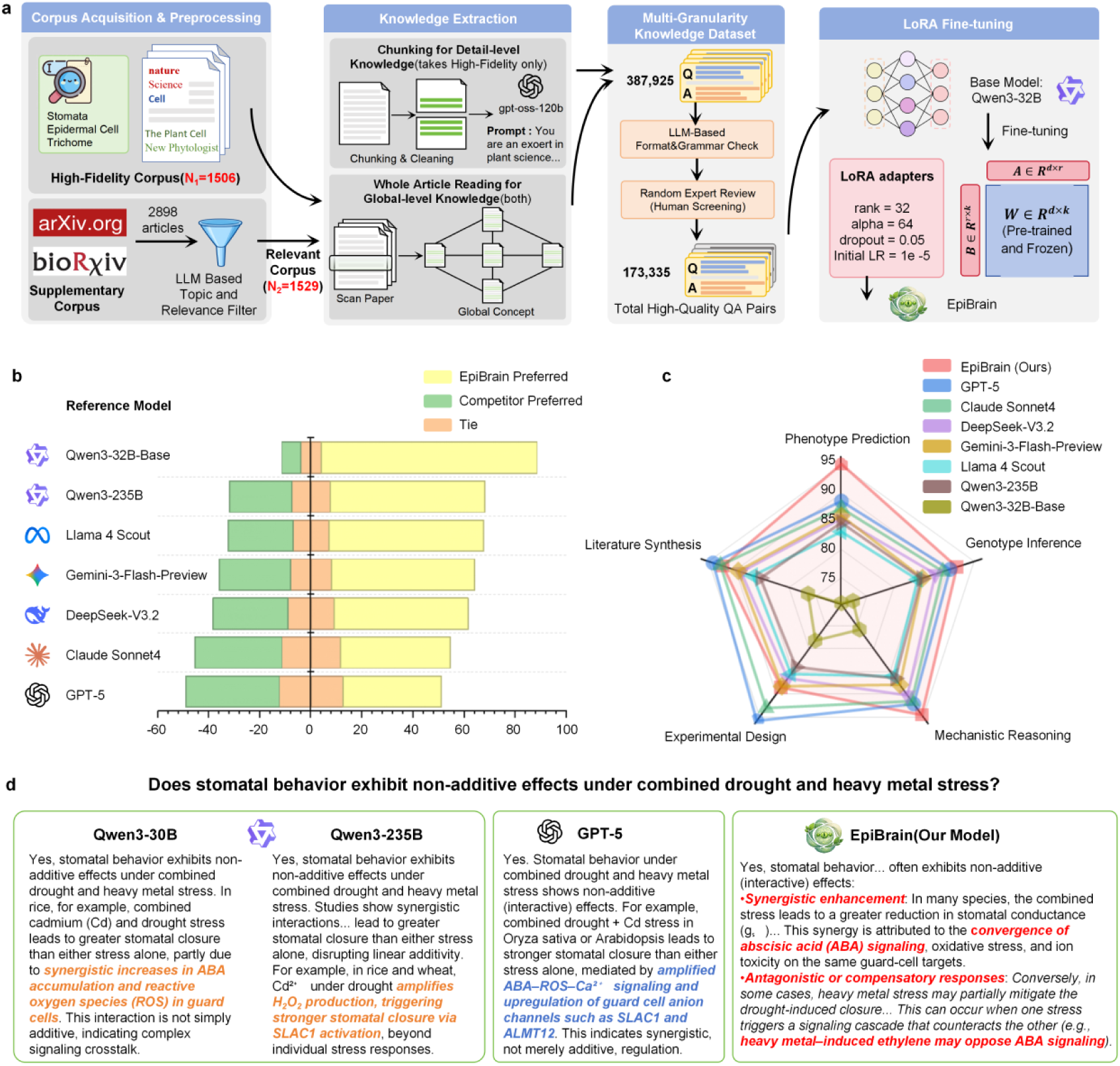
Construction workflow of the EpiBrain knowledge reasoning module and comparative performance of EpiBrain against general-purpose large language models. **a**, Corpus construction and model fine-tuning architecture of EpiBrain. The comprehensive workflow integrates four core modules: literature collection, knowledge extraction, data quality control comprising large model validation and expert blind review, and PEFT of Qwen3-32B utilizing LoRA technology. **b**, Benchmark evaluation across 30 stomatal biology questions. The divergent bar chart illustrates the proportions of win, tie, and loss preferences for EpiBrain compared with the foundation model and general-purpose large models when processing domain-specific instructions. **c**, Quantitative benchmarking of multidimensional professional capabilities. The radar chart compares the scoring performance of the evaluated models across phenotype prediction, genotype inference, mechanistic reasoning, experimental design, and literature synthesis. **d**, Example responses to the query regarding whether stomatal behavior exhibits non-additive effects under combined drought and heavy metal stress.

During construction, scientific knowledge graphs and LLMs were initially utilized to execute strict topic relevance filtering, screening high-quality content from 1,506 non-open-access articles published in the authoritative journals such as Nature, Science, Cell, and The Plant Cell, alongside 2,898 open-access cutting-edge preprints from arXiv and bioRxiv. Subsequently, in the knowledge extraction phase, fine-grained text chunking was adopted to conduct full-text reading and deep semantic parsing of the literature corpus. Through syntax-level cross-validation by large models combined with double-blind review by domain experts, an instruction fine-tuning dataset containing 173,335 high-quality question-answer pairs was constructed. Based on this foundation, parameter-efficient fine-tuning (PEFT) via LoRA technology was conducted on the Qwen3-32B foundation model. This approach retained the general semantic comprehension capabilities of the model while integrating exclusive domain knowledge related to plant epidermal features, particularly stomata, thereby establishing the EpiBrain reasoning module.

To evaluate the reliability of EpiBrain, we conducted multidimensional model alignment and human preference benchmarking. In a double-blind evaluation led by domain experts (Fig. 4b), EpiBrain was compared with frontier general LLMs, including GPT-5, Claude Sonnet 4, DeepSeek-V3.2, and Qwen3-235B, as well as the non-fine-tuned foundation model (Qwen3-32B-Base). The results indicated that the domain-fine-tuned EpiBrain exhibited a significant domain advantage in the human preference rate. Compared to the original foundation model, EpiBrain demonstrated substantial superiority with a preference rate of 84.5% (the foundation model was merely 7.5%). In comparisons with general models, when facing the hundred-billion-parameter-scale Qwen3-235B and Claude Sonnet 4, the preference rate of EpiBrain increased by 35.0% and 9.0%, respectively. Notably, EpiBrain maintained a competitive edge with a preference rate of 38.5% compared to 36.5% for GPT-5, which possesses the strongest comprehensive capabilities, with the remaining 25% of cases resulting in a tie.

Furthermore, through double-blind evaluation, we analyzed and compared the response capabilities of the 8 models across phenotypic prediction, genotypic inference, mechanistic reasoning, experimental design, and literature synthesis (Fig. 4c). In dimensions reflecting general scientific literacy, recently developed general models, benefiting from their ultra-large-scale pre-training corpora. It exhibited superior performance compared to EpiBrain in the indices of experimental design and literature synthesis, specifically GPT-5 and Claude Sonnet 4. However, for specialized questions highly dependent on vertical-domain prior knowledge, EpiBrain outperformed existing large models, most notably in phenotype prediction, where it scored 95.2, significantly leading GPT-5 at 88.6. These results demonstrate that the vertical-domain fine-tuning strategy can overcome the limitations of general models in specialized scientific fields. We took the query of whether stomatal traits exhibit non-additive effects under combined drought and heavy metal stress as an example (Fig. 4d). General models such as GPT-5 and Qwen3-235B could retrieve the general concept of non-additive characteristics, but their deductive logic often remained at the description of surface-level concepts, rarely engaging in logical reasoning on the underlying molecular regulatory mechanisms. Conversely, EpiBrain reasoning elucidated synergistic enhancement, specifically the convergence effect based on the abscisic acid (ABA) signaling pathway under the combined stress, characterizing it as an antagonistic or compensatory response where the heavy metal-induced ethylene pathway consequently blocks the ABA signal. Thus, EpiBrain possesses comprehensive and professional knowledge in fields related to the plant epidermis, thereby potentially assisting researchers in accelerating scientific discovery.

### EpiReasoner-assisted genetic mapping and mechanistic elucidation of tomato stomatal traits

Furthermore, we explored the research application of EpiReasoner for quantifying numbers of stomata within a natural population of tomato (Fig. 5). Initially, EpiVision was utilized to perform automated segmentation on leaf epidermal microscopic samples derived from 170 tomato accessions, and stomatal density data were acquired based on the segmentation masks (Fig. 5b). Subsequently, GWAS was conducted to link stomatal density with high-quality genetic markers. Following stringent filtration, 1,430,699 high-quality single nucleotide polymorphism (SNP) loci were retained from approximately 8.66 million raw loci. Under a linear mixed model with strict corrections for population structure and kinship (genomic inflation factor *λ* ≈ 1.02), the Manhattan plot (Fig. 5c) illustrated eight significantly associated QTLs distributed across the tomato genome (*P* < 10^-5^). Among these, a major QTL was precisely mapped to a candidate interval on chromosome 8 (3,213,579 to 3,215,218 base pairs). Integrated with local linkage disequilibrium decay analysis (Fig. 5d), the peak SNP within this interval (*P* = 3.35 × 10^-6^) was anchored to the intronic region of the gene *LOC101244792*. Annotation information indicated that this candidate gene encodes an F-box family protein, which is designated herein as *SKIP19L*. Association analysis between genotypes and phenotypes demonstrated that samples carrying the heterozygous allele at this locus exhibited significantly elevated stomatal density (*P* = 0.0018). Furthermore, the heterozygosity rate of this locus reached 82.5% in the wild ancestor *Solanum pimpinellifolium*, whereas it significantly decreased to 5.5% in modern cultivated tomato (*S. lycopersicum*) (Fig. 5f), implying that the *SKIP19L* locus likely underwent intense artificial selection and experienced a severe domestication bottleneck, consistent with the broader genomic history of tomato breeding^35^. This drift in allele frequency is highly consistent with the domestication trend of stomatal density. Wild tomatoes require higher stomatal densities to cope with variable environmental stresses, whereas under the conditions of controlled environment agriculture (CEA) for cultivated tomatoes, maintain a lower stomatal density is a valuable trait for reducing transpiration and enhancing water use efficiency alongside fruit yield^36,37^.

**Fig. 5.**
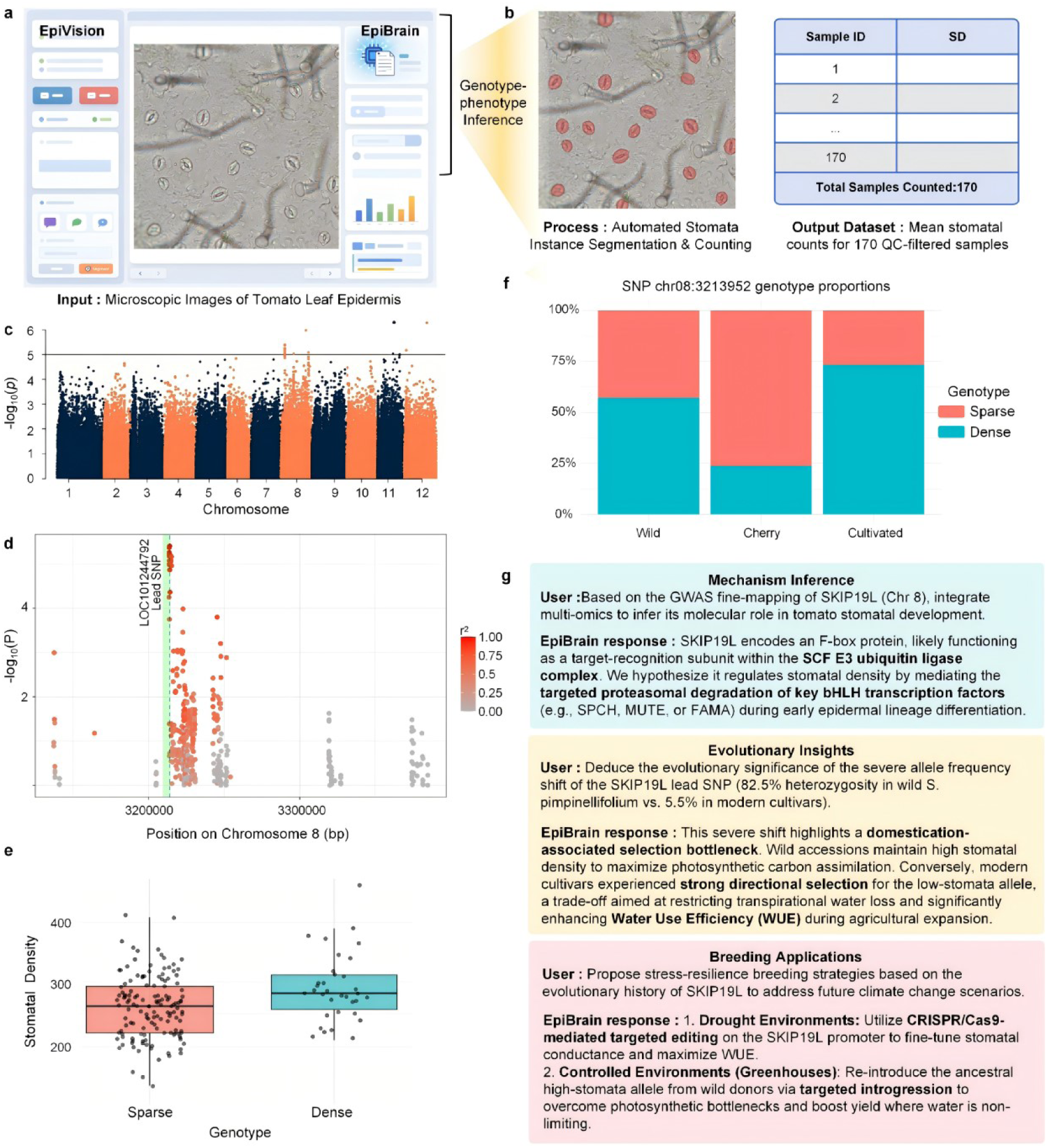
EpiReasoner-assisted genetic mapping and mechanistic elucidation of tomato stomatal density. **a**, Automated extraction of stomatal density phenotypic data based on EpiVision. **b**, Stomatal density across 170 tomato natural population samples generated through batch processing by EpiVision. **c**, Manhattan plot of the genome-wide association study for stomatal density. **d**, Fine mapping and local linkage disequilibrium analysis of the major quantitative trait locus interval on chromosome 8. The peak SNP is precisely anchored to the intronic region of the candidate gene *SKIP19L*, with scatter colors encoding the r^2^ values between surrounding variant loci and the peak SNP. **e**, Genotype-phenotype association analysis of the peak locus. The box plot confirms significant quantitative variances in stomatal density among the germplasm populations carrying the sparse and dense alleles. **f**, Allele frequency drift of the *SKIP19L* locus during tomato domestication. The comparison illustrates the frequency distribution of this variant between wild ancestors and modern cultivated accessions, revealing the intense artificial selection it experienced. **g**, Biological hypotheses driven by EpiBrain.

In traditional forward genetics research, the progression from obtaining mapping results to formulating mechanistic reasoning hypotheses typically requires researchers to possess substantial research experience and conduct time-consuming literature reviews. Here, we introduce the EpiBrain LLM module to accelerate this process of scientific discovery. The candidate gene *SKIP19L* and its domestication selection features were input into the EpiBrain module to assist in deducing the potential molecular regulatory network (Fig. 5g). Based on relevant knowledge graphs, EpiBrain proposed the following hypothesis: SKIP19L may function as a target recognition subunit of the SKP1-CULLIN-F-BOX (SCF) ubiquitin ligase complex, regulating stomatal density by targeting key basic helix-loop-helix (bHLH) transcription factors in stomatal development (such as SPCH, MUTE, or FAMA) for degradation. Concurrently, from the perspective of evolutionary adaptation, the system performed theoretical modeling on the trade-offs of this trait during domestication. Wild tomato species retain high densities to maximize photosynthetic efficiency, whereas cultivated species tend to reduce density under artificial selection to enhance water use efficiency. Finally, the model also proposed utilizing targeted gene editing to regulate this locus as a strategy to accelerate stress-resistance breeding.

## Discussion

EpiReasoner represents a significant advance in plant stomatal researches by directly coupling high-fidelity visual trait extraction with domain-specialized mechanistic reasoning, overcoming two persistent barriers in plant biology: the confinement of phenotypic analysis models for epidermal cells such as stomata to a limited number of species and imaging modalities, which results in weak generalization and transferability, and the absence of domain-specific reasoning models and knowledge bases.

Based on optical microscopy samples from monocotyledonous and dicotyledonous plants, we employed a transfer learning strategy to conduct targeted optimization of the high-level feature decoding network within SAM3. Comparative analysis indicates that the performance of this model in the segmentation and recognition tasks of stomata and pavement cells has surpassed that of current proprietary supervised models. Our result further corroborates that current general vision models, such as the SAM series, possess robust fundamental visual comprehension capabilities, enabling efficient adaptation to various downstream visual tasks via fine-tuning or prompt mechanisms. This trend has been similarly validated in other disciplines^38,39^, demonstrating that fine-tuning foundation models on dedicated datasets can effectively adapt general vision models to the unique structural distributions and textural features of microscopic biological materials. Therefore, fully leveraging foundation models and combining them with domain-specific samples for targeted adaptation constitutes a highly efficient pathway for advancing plant science.

Currently, phenotypic analyses of cells such as stomata remain predominantly confined to the statistical description of basic geometric features, including area, perimeter, and aspect ratio^40,41^, lacking a quantitative index system capable of characterizing the complex topological morphology and spatial structural patterns of cells. Consequently, building upon the achievement of precise segmentation, this study further proposes 19 indices for the quantitative characterization of the complex morphology of stomata and pavement cells, alongside 4 indices delineating cellular spatial distribution patterns, thereby providing a novel analytical methodology for the systematic resolution of the morphology and tissue structure of epidermal cells such as stomata. Stomatal index (SI) is frequently utilized to reflect the developmental plasticity and adaptive strategies of plants in response to environmental fluctuations, such as carbon dioxide concentration, water stress, and light conditions^42–44^. The SCVR proposed in this study exhibits greater sensitivity to deviations from the one-cell spacing rule compared with the Stomatal Index, enabling the effective differentiation of phenotypic characteristics among various tomato mutants, including spatial distribution patterns and density variations of stomata.

The integration of EpiBrain further assists researchers in conducting biological reasoning of phenotypes, as well as retrieving and validating biological hypotheses. In the analysis of tomato natural variation, GWAS identified the *SKIP19L* locus using the standardized stomatal density data generated by EpiVision automatically. The severe drift in its allele frequency from wild ancestors to cultivated accessions completely aligns with the domestication trend of reducing transpiration and enhancing water use efficiency under artificial selection. The hypothesis proposed by EpiBrain suggests that *SKIP19L* may act as an SCF ubiquitin ligase adapter targeting bHLH transcription factors such as *SPCH*, *MUTE*, or *FAMA* for degradation^45^. This integrates known stomatal lineage regulatory networks with the logic of evolutionary adaptive trade-offs, providing a testable strategy for CRISPR-mediated gene-editing breeding. This closed-loop phenotype-genotype workflow compresses the traditional process, which requires months of literature review and expert synthesis, into a reproducible streamlined intelligent pipeline.

Although EpiVision generalizes well, rare epidermal morphologies or novel imaging platforms may still require additional fine-tuning. While extensive, the knowledge base of EpiBrain is derived from current literature and will benefit from continuous database updating. Future extensions could incorporate multi-omics integration, three-dimensional epidermal reconstruction, and/or real-time field phenotyping. Nevertheless, by providing a unified, open framework spanning visual perception and knowledge reasoning, EpiReasoner establishes a novel paradigm for accelerating mechanistic discovery and translational breeding for key crops such as tomato and wheat.

## Methods

### Construction and preprocessing of the dataset

During the fine-tuning phase of the vision model, this study acquired and constructed an in-house training dataset utilizing bright-field microscopy. This dataset encompasses 170 samples of the dicotyledonous plant tomato (*Solanum lycopersicum*) and 210 samples of the monocotyledonous plant wheat (*Triticum aestivum*), prepared via the epidermal peeling method. Considering the variations in physical scale and observation targets among microscopic samples, to adapt to network computation, the acquired raw samples were subjected to non-overlapping cropping, generating a total of 530,470 standardized patches with a resolution of 512 × 512 pixels. Subsequently, a representative subset of patches was annotated utilizing the ISAT annotation software. To ensure the learning capacity of the model for pavement cells with complex morphologies, this study annotated 407 tomato patches and 58 wheat patches. Within this annotated subset, a total of 5,419 pavement cells, 1,471 stomatal complexes, and 629 stomatal pores were extracted from the tomato samples; correspondingly, 1,780 pavement cells, 426 stomatal complexes, and 51 stomatal pores were extracted from the wheat samples.

To validate the cross-domain generalization capability of EpiVision, this study further integrated the in-house data with 13 open-source databases to construct a large-scale heterogeneous validation set, EpiDataset. This dataset encompasses three sample preparation techniques, including epidermal peeling, impression methods, and in-situ imaging, alongside three imaging modalities, namely bright-field, SEM, and DIC. During the model inference phase, all patches were subjected to standardized normalization based on the mean and standard deviation of ImageNet. Furthermore, to eliminate the interference of resolution discrepancies across cross-source samples on feature extraction, the raw samples from the open-source databases were similarly subjected to non-overlapping cropping at a resolution of 512 × 512 pixels. Upon integration of these standardized open-source data with the in-house data, they collectively constituted the EpiDataset, comprising 993,833 patches of 512 × 512 pixels. Prior to extracting deep features and performing dimensionality reduction visualization via UMAP, to prevent majority-class samples from dominating the feature space, a rigorous stratified random sampling strategy was implemented. Based on each attribute intersection category delineated by UpSet analysis, a maximum sampling threshold of 300 patches was established; for long-tail categories with sample sizes below this threshold, all instances were fully retained.

### Construction and fine-tuning of the EpiVision model

The EpiVision model utilizes the foundation model SAM3 as its architecture. Although SAM3 demonstrates robust feature extraction capabilities in natural scene scenarios, it lacks the semantic representation capacity for botanical concepts such as plant stomata or pavement cells. Regarding the model fine-tuning strategy, the core weights of the underlying visual feature extractor were frozen, and a transfer learning strategy was adopted to conduct targeted fine-tuning of the high-level feature decoding network. Specifically, the weight updates of the DETR decoder were enabled, and the perception head, bounding box head, and mask head were targeted and fine-tuned.

During the data preprocessing phase, a dynamic interactive prompt simulation mechanism encompassing random point and bounding box sampling was introduced, and the input samples were uniformly processed to a resolution of 1008 × 1008. Model training was deployed on the PyTorch distributed data parallel framework, supported by a high-performance computing cluster equipped with an 8 × NVIDIA H800 (80 GB) GPU node interconnected via NVLink. The process fully enabled Bfloat16 mixed-precision computation, employing the AdamW optimizer, an inverse square root scheduler, and a layer-wise learning rate decay strategy for specific modules to ensure the convergence stability of the deep feature network. Regarding the optimization objective, the network executed sample assignment via an improved bipartite Hungarian matching algorithm and applied a multitask joint loss function deeply coupling bounding box regression, open-vocabulary semantic classification, and a highly weighted pixel-level segmentation loss, thereby strictly constraining the fitting precision of the model for complex cellular mosaic boundaries.

### Model performance evaluation

To comprehensively and objectively evaluate the performance of the EpiVision model in the multi-cell instance segmentation task of the plant leaf epidermis, this study constructed a multidimensional comprehensive evaluation system and conducted rigorous quantitative comparisons with current mainstream baseline models, including Mask R-CNN, OneFormer, and YOLO26. To ensure evaluation fairness, all comparative models adopted identical training settings and test set partitions.

During the inference phase, all input samples were uniformly resized to a resolution of 1008 × 1008 and subjected to standardization. To eliminate redundant predictions and enhance mask quality, the preliminary prediction results output by the model required a rigorous post-processing pipeline. Initially, a confidence threshold of 0.5 and non-maximum suppression (NMS) with a threshold of 0.5 were applied for instance screening. Subsequently, morphological opening and closing operations utilizing a 3 × 3 convolutional kernel were employed to smooth cellular boundaries, and minute noise regions with an area of less than 16 pixels were eliminated.

For the single-cell instance segmentation task, this study adopted precision, recall, F1-score, Dice coefficient, alongside mean average precision (mAP) and mean average recall (mAR) as foundational evaluation metrics (fundamental mathematical definitions are detailed in Supplementary Table 2). These metrics are primarily based on pixel-level or region-level overlap, effectively measuring overall segmentation accuracy.

Given that epidermal cells frequently exhibit a dense, jigsaw-like mosaic arrangement, conventional metrics struggle to acutely capture the phenomena of over-segmentation (a single cell erroneously segmented into multiple instances) or under-segmentation (multiple cells erroneously merged into a single instance). Therefore, this study introduced the AJI to more strictly evaluate the overall integrity of instance structures. By matching each ground-truth cell instance with its optimal predicted instance and accumulating the intersection over union (IoU) of all matched instances, while simultaneously incorporating unmatched predicted instances (false positives) into a penalty term, the AJI effectively reflects the instance separation and matching capabilities of the model in dense scenarios.

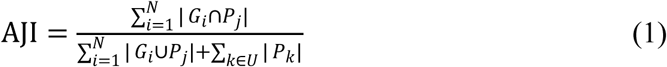

where *G*_*i*_ represents the *i*-th ground-truth cell instance, *P_j_*is the predicted instance exhibiting the maximum IoU with it, and *U* denotes the set of predicted instances unmatched to any ground-truth targets, specifically the false positive instances.

Furthermore, addressing the highly irregular, lobed topological features of pavement cell margins, conventional evaluation metrics based on regional overlap are easily dominated by large areas of correct internal cellular segmentation, rendering them insufficiently sensitive to reflect the fitting capacity at microscopic boundaries. Consequently, this study further introduced geometric evaluation metrics focusing on boundary quality: BoundIoU, Normalized Surface Distance (NSD), and 95% Hausdorff Distance (HD95).

BoundIoU specifically quantifies the concordance of boundary contours within a given dilation threshold range. For lobed cells, a higher BoundIoU value indicates that the predicted contours of the model at complex concave and convex margins adhere more closely to the authentic morphology, rather than merely relying on overall area overlap.

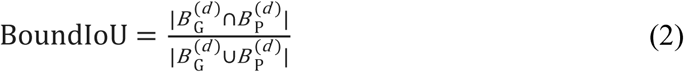

where 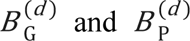 denote the dilated pixel sets of the ground-truth and predicted mask boundaries within the Euclidean distance threshold *d*.

The NSD is utilized to evaluate the proportion of mutual coverage between predicted and ground-truth boundaries within a specified tolerance (set to 2 pixels in this study). This metric introduces a reasonable tolerance for boundary uncertainty, rendering it particularly suitable for minor annotation ambiguity or imaging noise present in microscopic samples. A higher NSD value signifies a stronger capacity of the model to capture minute boundary details.

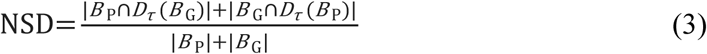

where *B*_P_ and *B*_G_ denote the boundary pixel sets of the predicted and ground-truth masks, respectively, and *D_τ_*(·) represents the distance transform function within the tolerance distance range τ.

The Hausdorff Distance is employed to measure the maximum degree of mismatch between two boundary sets. To eliminate the excessive interference of extremely isolated outlier noise on the overall metric, this study adopted its 95th percentile (HD95, with the normalization coefficient set to 0.2). A lower HD95 value indicates a smaller maximum local deviation on the predicted boundaries of the model, effectively revealing the robustness of the model when processing extremely irregular geometric morphologies such as lobe tips or invaginations.

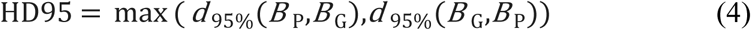

where *d* _95%_(*X*,*Y*) represents the 95th percentile of the shortest Euclidean distances from all points in set *X* to set *Y*.

### Algorithms for phenotypic quantification and topological network calculation

Following the acquisition of instance segmentation masks, the algorithm initially eliminates absolute size measurement errors caused by varying microscopic field-of-view scaling through physical scale calibration. Based on this, the present study establishes a phenotypic quantification methodology targeting the complex geometric morphologies of stomatal complexes and pavement cells, alongside their spatial distribution relationships.

For stomatal complexes, the system extracts the stomatal area (*A*_stoma_), perimeter (*P*_stoma_), and Feret’s diameter (*D*_F_) based on the structural contours. Utilizing Principal Component Analysis (PCA) to resolve the eigenvalues and eigenvectors of the coordinate covariance matrix, the algorithm quantifies the major axis length (*L_major_*), minor axis length (*L*_minor_), and spatial deflection angle (*θ*) of the stomata.

Integrating the geometric relationship between the outer contour of the stomata and the internal pore, the algorithm further extracts the pore area (*A*_pore_), the major axis length of the pore (*L*_pore_major_), and the minor axis length (*L*_pore_minor_). To evaluate the degree of stomatal opening, the stomatal aperture index (*I*_aperture_) and pore area fraction (*R*_pore_) are calculated according to the following formulas:

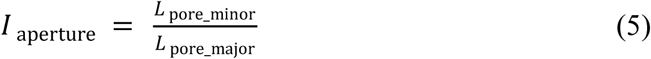

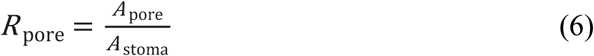

Distinct from the relatively regular elliptical structure of stomata, to address the complex lobed contours of pavement cells, this study utilizes Circularity (*C*), Solidity (*S*), and Rectangularity (*R*) to quantify their overall morphological complexity, defined as follows:

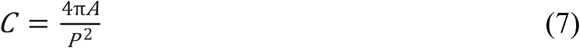

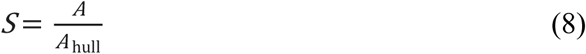

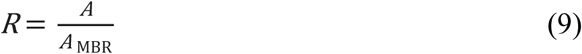

where *A* represents the cellular pixel area, *P*is the contour perimeter, *A*_hull_ denotes the area of the corresponding convex hull, and *A*_MBR_ is the area of the minimum bounding rectangle. To address specific lobed morphological structures, the algorithm achieves automated determination of local lobe features by locating the local curvature (*κ*) maxima (i.e., lobe tip points, *p*_tip_) and convexity defects (i.e., lobe invagination points, *p*_defec*t*_) along the contour curve, integrated with the morphological skeleton and network betweenness centrality.The neck width *W*_neck_ is defined as the shortest Euclidean distance between two opposing convexity defects, representing the width at the morphological bottleneck of the cell, namely:

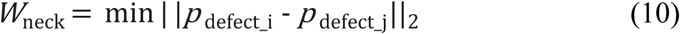

The lobe length (*L*_lobe_) is defined as the Euclidean distance from the lobe tip to the midpoint (*p*_midpoint_) of the line connecting its corresponding neck, namely:

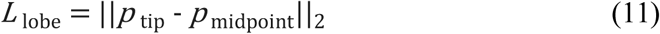

The protrusion depth (*D*_protrusion_) is defined as the vertical projection distance from the lobe tip to the neck baseline formed by the line connecting two adjacent convexity defects. Furthermore, based on the Euclidean distance transform from internal cellular pixels to the contour edge, the system computes a cellular thickness heatmap reflecting the internal expansion state, thereby inferring the physical and mechanical stress distribution during cellular deformation.

Following the extraction of the morphological features of stomata and pavement cells, this study further extracts the indices utilized to calculate stomatal abundance: stomatal density(SD) and SI.SD is defined as the absolute number of stomata per unit epidermal area, calculated as follows:

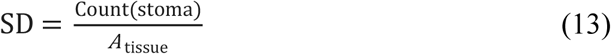

where the parameter Count(stoma) denotes the total number of stomata within the analyzed region, and *A*_tissue_ represents the corresponding absolute physical area.

The SI is calculated as follows:

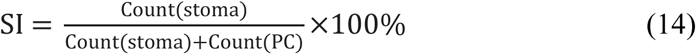

where the parameter Count(PC) represents the total number of extracted pavement cells within the identical region.

This study further constructs a spatial topological relationship model between stomata and pavement cells. Initially, based on the cellular topological adjacency graph defined by physical boundary contacts, the spatial connectivity network of stomata is reconstructed by extracting the geometric centroids of all stomata and applying Delaunay triangulation. To quantitatively validate the one-cell spacing rule in plant epidermal development, the algorithm executes a breadth-first search (BFS) on the adjacency graph composed of pavement cells to calculate the shortest topological path *D*_topo_(*s*_*i*_ + *s*_*j*_) connecting any two adjacent stomatal nodes *s_i_* and *s_j_*, representing the actual number of intervening pavement cells between them. Its calculation formula is:

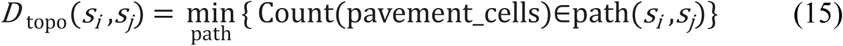

Based on this path value, the system conducts rigorous qualitative and quantitative classification of the stomatal distribution patterns within the population: *D*_topo_ = 0 indicates direct physical contact between adjacent stomata, which the system determines as a developmental cluster violation; *D*_topo_ = 1 represents adherence to the normal single-cell interval; whereas *D*_topo_ ≥ 2 is classified as a relaxed interval.

On this basis, this study establishes the SCVR index. This metric computes the occurrence intensity of all cluster violation events within a specified area of the epidermal tissue, defined as the proportion of adjacency relationships exhibiting direct physical contact (*D*_topo_ = 0) among all stomatal connectivity relationships. Its mathematical expression is:

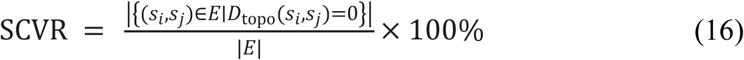

where the parameter *E* represents the complete set of stomatal adjacency edges constructed via Delaunay triangulation. The SCVR serves as a normalized population-level metric for the degree of spatial deviation from the one-cell spacing rule among stomata, ranging from 0% to 100%. A higher value indicates a more prevalent phenomenon of violating clustered stomata within the epidermal tissue.

Furthermore, in addition to focusing on stomatal distribution patterns, to delineate the topological relationships of the epidermal tissue under physical tension, this study introduces the Tricellular Junction Density (*ρ*_TCJ_) as a quantitative index. This study utilizes morphological gradients and dilation algorithms to calculate the physical boundaries of pavement cells, locating and extracting topological nodes where three or more pavement cells mutually intersect (designated as tricellular junctions (TCJs)). Subsequently, the junction density is acquired by calculating the ratio of the total number of tricellular junctions within the analyzed region to the total area of the epidermal tissue. The calculation formula is:

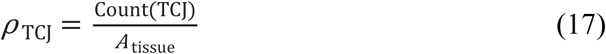

where the parameter Count(TCJ) denotes the total number of tricellular junctions detected within the region, and *A*_tissue_ represents the area of that region. A higher ρ_TCJ_ value indicates a denser concentration of pavement cell intersection points per unit area.

### Corpus construction and fine-tuning of the EpiBrain knowledge reasoning module

The EpiBrain reasoning module was constructed based on the open-source large language model Qwen3-32B, with the specific pipeline detailed as follows:

During the data acquisition and preprocessing phase, full-text articles from 1,506 publications regarding leaf epidermal features, such as stomata and pavement cells, from journals such as *Nature*, *Science*, *Cell*, *The Plant Cell*, alongside 1,529 recent manuscripts acquired from the arXiv and bioRxiv preprint platforms, were collected and subjected to manual screening.

In the automated knowledge extraction phase, a 120-billion-parameter open-source large model (gpt-oss:120b) was deployed locally as the data generation model, and differentiated extraction strategies were formulated for data sources exhibiting varying knowledge densities. For core journal literature, the system executed a more refined data cleaning process. Initially, the Easy Dataset v1.6.1 tool^46^ was utilized to segment the literature into contextual blocks of 2,500 to 4,000 characters, and the large model was invoked to filter out residual non-academic noise, such as isolated symbols and supplementary material annotations, thereby retaining the descriptive text. Building upon this foundation, a customized prompt generator was employed to guide the model in generating questions investigating causal relationships centered on concepts including stomata and pavement cells. Concurrently, the system restricted the model from generating metalanguage such as “according to this paper” or “as shown in the figure” in its responses, thereby extracting context-independent replies and mitigating the risk of the model overfitting to specific textual structures during subsequent fine-tuning. For literature such as preprints, PyMuPDF was utilized to parse the text, and regular expressions were applied to filter redundant information including citation markers, URLs, and acknowledgments. Subsequently, by establishing domain expert role prompts, the model was constrained to generate question-answering data encompassing experimental design, data interpretation, and critical analysis based on the cleaned text. Multiple rounds of interactive data were iteratively generated and ultimately encapsulated into the standardized ShareGPT format to construct a global macroscopic conceptual corpus.

All generated question-answer pairs were subjected to a multidimensional quality evaluation pipeline. During the automated filtration phase, an evaluation agent based on large language models (LLM-as-a-judge) was introduced to systematically score the data across 4 dimensions: 1) question quality (intent clarity and academic rationality); 2) answer quality (logical coherence and accuracy); 3) text relevance (the anchoring degree of the question-answer content to the original text); and 4) global consistency (the closed-loop integrity of the reasoning logic). Based on the output evaluation scores, the system eliminated data exhibiting factual errors, logical flaws, or low relevance. Following the automated cleaning, supplemented by expert blind sampling review, an instruction fine-tuning dataset containing 173,335 high-quality question-answer pairs was ultimately constructed.

In the supervised fine-tuning phase, PEFT was conducted on the Qwen3-32B foundation model utilizing LoRA technology within the consistent hardware environment. LoRA adapters were applied to all linear layers of the model to enhance the fitting capacity for domain features. The relevant hyperparameters were established as follows: a rank of 32, a scaling factor of 64, and a dropout rate of 0.05 to mitigate overfitting. The training process enabled the DeepSpeed ZeRO-2 strategy, Bfloat16 mixed precision, and gradient checkpointing technologies to optimize the memory efficiency of distributed training. The maximum context sequence length of the model was set to 4096 tokens, with a global batch size of 96. The AdamW optimizer was selected in combination with a cosine annealing learning rate scheduler, utilizing an initial learning rate of 1e-5, a warmup ratio of 3%, and a weight decay of 0.1. Based on the 173,335 customized corpus entries, the model completed two epochs of iterative training, ultimately yielding the EpiBrain reasoning module.

### Model inference and benchmark testing pipeline

To objectively evaluate the domain knowledge reasoning capabilities of the respective models on a unified scale, this study developed an automated benchmarking pipeline based on Python and the vLLM inference engine, deployed on the identical computing nodes. Regarding the construction of the test set, we designed a set of 30 specialized questions encompassing biotic stress responses, abiotic stress responses, environmental signaling, and molecular mechanisms. These questions deliberately eschewed basic definitional queries, focusing instead on assessing the capacity of the models for in-depth mechanistic dissection of complex signaling pathway integration, cross-species regulatory discrepancies, and non-additive effects. Concerning prompt engineering and inference configurations, to eliminate biases introduced by the inherent response styles of different LLMs and ensure the fairness of horizontal comparisons, we adopted a style alignment strategy. For all general-purpose LLMs and foundation models that lacked domain-specific fine-tuning, we mandated the use of precise, direct, and high-information-density academic expressions within the system prompts. Combined with few-shot prompting, the models were constrained to output conclusions directly, strictly prohibiting the generation of verbose and tedious background filler. For EpiBrain, the domain expert role prompt, which was entirely consistent with its fine-tuning phase, was employed. Regarding the establishment of inference hyperparameters, the generation temperature of all models was strictly set to 0, and the random seed was set to 42, thereby eliminating the randomness in the autoregressive decoding process to ensure the reproducibility and scientific rigor of the evaluation results. The responses generated by the models were automatically collected and stored in a structured matrix, subsequently advancing to the downstream expert double-blind and performance evaluation scoring pipeline.

### Expert double-blind and performance evaluation

Following the acquisition of the standardized inference outputs from all evaluated models, a rigorous expert double-blind scoring mechanism and evaluation pipeline were executed to systematically assess the quality of the generated content. In terms of double-blind qualitative evaluation, all text generated by the models was stripped of identifiers and submitted to senior domestic and international experts with backgrounds in botany and developmental biology for independent review, ultimately yielding 8 survey results. In pairwise comparisons, experts were required to adjudicate the responses based on scientific accuracy, logical coherence, and the substantiation of mechanistic evidence, subsequently classifying them as a win or a tie. Concurrently, the evaluation system introduced a strict single-veto mechanism for factual errors. Any response containing erroneous pathway splicing, fictitious gene nomenclature, or contradictions of known physiological common sense was directly recorded as an error to evaluate the probability of the models generating hallucinations within the vertical domain. Regarding fine-grained scoring, with reference to the specific scientific research scenarios of botanical research, we designed a centesimal scoring standard encompassing five dimensions: literature synthesis evaluated the capacity of the model to extract knowledge from redundant information; experimental design assessed the logicality of the model in proposing reasonable validation schemes for specific genes or phenotypes; phenotype prediction evaluated the accuracy of the model in deducing the macroscopic phenotypes of crops and cells, such as stomata, based on specific mutations or environmental stress conditions; genotype inference assessed the capability of the model to deduce potential gene functions through specific developmental defect phenotypes; and molecular mechanism reasoning evaluated the professional depth of the model in deconstructing and recombining complex signaling cascade networks.

### Genome-wide association studies (GWAS)

This study utilized EpiVision to perform automated segmentation on bright-field epidermal microscopic samples derived from a tomato germplasm population of 170 accessions, and extracted the stomatal density data for all samples. Regarding genotypic data, the raw variation set comprised 8.66 million SNP loci. Based on the established quality control criteria, specifying a minor allele frequency (MAF) ≥ 0.05, an SNP missing rate ≤ 0.05, and a Hardy-Weinberg equilibrium *P* ≥ 10^-6^, 1,430,699 high-quality SNPs and 170 qualified samples were ultimately retained for subsequent analysis.

The GWAS was executed utilizing PLINK v1.90 software^47^. To rigorously correct for false positives induced by population stratification, the analysis was based on a unified mixed-model framework^48^, incorporating the first three principal components calculated via PCA as covariates to correct for false positives induced by population stratification. The significance threshold was established at a *P* < 10^-5^. For the screened major QTL, integrated with the local LD decay profile, candidate gene annotation was conducted utilizing the Bedtools closest tool with a distance threshold of 10 kilobases based on the NCBI SL3.0 tomato reference genome. In accordance with allele tracing, the alterations in allele frequency and heterozygosity of the critical variant loci from the wild ancestor, S.*pimpinellifolium*, to the modern cultivated population were statistically analyzed.

## Acknowledgement

This work was supported by the National Natural Science Foundation of China (32572179)

## Data Availability

Data will be made available on request.

## Supplementary Information

**Supplementary Table 1.** Summary of publicly available open-source stomatal and leaf epidermis image datasets used in this study.

**Supplementary Table 2.** Evaluation metrics for assessing instance segmentation performance of stomatal and pavement cells.

**Supplementary Table 3.** Phenotypic traits of stomatal complexes and pavement cells with their calculation methods.

## References

1. Liu, S., Jobert, F., Rahneshan, Z., Doyle, S. M. & Robert, S. Solving the Puzzle of Shape Regulation in Plant Epidermal Pavement Cells. Annu. Rev. Plant Biol. 72, 525–550 (2021).

2. Kheibarshekan Asl, L., et al. Model-Based Analysis of Arabidopsis Leaf Epidermal Cells Reveals Distinct Division and Expansion Patterns for Pavement and Guard Cells. Plant Physiology 156, 2172–2183 (2011).

3. Melotto, M., Underwood, W. & He, S. Y. Role of Stomata in Plant Innate Immunity and Foliar Bacterial Diseases. Annu. Rev. Phytopathol. 46, 101–122 (2008).

4. Lawson, T. & Matthews, J. Guard Cell Metabolism and Stomatal Function. Annu. Rev. Plant Biol. 71, 273–302 (2020).

5. Li, S. et al. LeafNet: a tool for segmenting and quantifying stomata and pavement cells. Plant Cell 34, 1171–1188 (2022).

6. Fetter, K. C., Eberhardt, S., Barclay, R. S., Wing, S. & Keller, S. R. StomataCounter: a neural network for automatic stomata identification and counting. New Phytologist 223, 1671–1681 (2019).

7. Sai, N. et al. StomaAI: an efficient and user-friendly tool for measurement of stomatal pores and density using deep computer vision. New Phytol. 238, 904–915 (2023).

8. Wang, J., Renninger, H. J., Ma, Q. & Jin, S. Measuring stomatal and guard cell metrics for plant physiology and growth using StoManager1. Plant Physiology 195, 378–394 (2024).

9. Hills, A., Chen, Z.-H., Amtmann, A., Blatt, M. R. & Lew, V. L. OnGuard, a Computational Platform for Quantitative Kinetic Modeling of Guard Cell Physiology. Plant Physiol. 159, 1026–1042 (2012).

10. Wang, Y. et al. Unexpected Connections between Humidity and Ion Transport Discovered Using a Model to Bridge Guard Cell-to-Leaf Scales. Plant Cell 29, 2921–2939 (2017).

11. Sun, Z. et al. StomataTracker: Revealing circadian rhythms of wheat stomata with in-situ video and deep learning. Computers and Electronics in Agriculture 212, 108120 (2023).

12. Vőfély, R. V., Gallagher, J., Pisano, G. D., Bartlett, M. & Braybrook, S. A. Of puzzles and pavements: a quantitative exploration of leaf epidermal cell shape. New Phytologist 221, 540–552 (2019).

13. Nowak, J. et al. A network-based framework for shape analysis enables accurate characterization of leaf epidermal cells. Nat Commun 12, 458 (2021).

14. Ma, J. et al. The multimodality cell segmentation challenge: toward universal solutions. Nat Methods 21, 1103–1113 (2024).

15. Archit, A. et al. Segment Anything for Microscopy. Nat Methods 10.1038/s41592-024-02580-4 (2025) doi:10.1038/s41592-024-02580-4.

16. Greenwald, N. F. et al. Whole-cell segmentation of tissue images with human-level performance using large-scale data annotation and deep learning. Nat Biotechnol 40, 555–565 (2022).

17. Stringer, C., Wang, T., Michaelos, M. & Pachitariu, M. Cellpose: a generalist algorithm for cellular segmentation. Nat Methods 18, 100–106 (2021).

18. Van Doorselaer, L., Verboven, P. & Nicolai, B. Automatic 3D cell segmentation of fruit parenchyma tissue from X-ray micro CT images using deep learning. Plant Methods 20, 12 (2024).

19. Zheng, Y. et al. Large language models for scientific discovery in molecular property prediction. Nat Mach Intell 7, 437–447 (2025).

20. Chen, S. F. LLM-assisted systematic review of large language models in clinical medicine. Nat. Med.

21. Yu, H. et al. PlantScience.ai: An LLM-powered virtual scientist for plant science. Molecular Plant 19, 1117–1123 (2026).

22. Zhang, R. et al. PlantGPT: An Arabidopsis-Based Intelligent Agent that Answers Questions about Plant Functional Genomics. Advanced Science e03926 (2025) doi:10.1002/advs.202503926.

23. Carion, N. et al. SAM 3: Segment Anything with Concepts.

24. Yang, A. et al. Qwen3 Technical Report. Preprint at 10.48550/arXiv.2505.09388 (2025).

25. Hu, E. et al. LORA: LOW-RANK ADAPTATION OF LARGE LAN- GUAGE MODELS. (2022).

26. He, K., Gkioxari, G., Dollár, P. & Girshick, R. Mask R-CNN. Preprint at 10.48550/arXiv.1703.06870 (2018).

27. Jain, J. et al. OneFormer: One Transformer to Rule Universal Image Segmentation. in 2023 IEEE/CVF Conference on Computer Vision and Pattern Recognition (CVPR) 2989–2998 (IEEE, Vancouver, BC, Canada, 2023). doi:10.1109/CVPR52729.2023.00292.

25. Sapkota, R., Cheppally, R. H., Sharda, A. & Karkee, M. YOLO26: Key Architectural Enhancements and Performance Benchmarking for Real-Time Object Detection. Preprint at 10.48550/arXiv.2509.25164 (2026).

29. Liang, X. et al. StomataScorer: a portable and high-throughput leaf stomata trait scorer combined with deep learning and an improved CV model. Plant Biotechnology Journal 20, 577–591 (2022).

30. McInnes, L., Healy, J. & Melville, J. UMAP: Uniform Manifold Approximation and Projection for Dimension Reduction. Preprint at 10.48550/arXiv.1802.03426 (2020).

31. Von Groll, U., Berger, D. & Altmann, T. The Subtilisin-Like Serine Protease SDD1 Mediates Cell-to-Cell Signaling during Arabidopsis Stomatal Development. Plant Cell 14, 1527–1539 (2002).

32. Jalakas, P., Tulva, I., Bērziņa, N. M. & Hõrak, H. Stomatal patterning is differently regulated in adaxial and abaxial epidermis in Arabidopsis. Journal of Experimental Botany 75, 6476–6488 (2024).

33. Geisler, M., Nadeau, J. & Sack, F. D. Oriented Asymmetric Divisions That Generate the Stomatal Spacing Pattern in Arabidopsis Are Disrupted by the too many mouths Mutation.

33. Yang, M. & Sack’, F. D. The too many mouths and four lips Mutations Affect Stomatal Pnoduction in Arabidopsis.

35. Lin, T. et al. Genomic analyses provide insights into the history of tomato breeding. Nat Genet 46, 1220–1226 (2014).

36. Muir, C. D., Pease, J. B. & Moyle, L. C. Quantitative Genetic Analysis Indicates Natural Selection on Leaf Phenotypes Across Wild Tomato Species (Solanum sect. Lycopersicon; Solanaceae). Genetics 198, 1629–1643 (2014).

37. Ganie, S. A. Unravelling the physiological and anatomical basis of divergent adaptations in cultivated and wild tomatoes.

38. Israel, U. et al. CellSAM: A Foundation Model for Cell Segmentation. Preprint at 10.1101/2023.11.17.567630 (2023).

39. Ma, J. et al. Segment anything in medical images. Nat Commun 15, 654 (2024).

40. Xie, J. et al. Optical topometry and machine learning to rapidly phenotype stomatal patterning traits for maize QTL mapping. Plant Physiology 187, 1462–1480 (2021).

41. Zhang, C. et al. Analysis of stomatal characteristics of maize hybrids and their parental inbred lines during critical reproductive periods. Front. Plant Sci. 15, 1442686 (2025).

42. Royer, D. L. Stomatal density and stomatal index as indicators of paleoatmospheric CO2 concentration. Review of Palaeobotany and Palynology 114, 1–28 (2001).

43. Qi, X. & Torii, K. U. Hormonal and environmental signals guiding stomatal development. BMC Biol 16, 21 (2018).

44. Poorter, H., Pons, T. L. & Reichgelt, T. Stomatal Density and Index Are More Responsive to Light Intensity than to [CO2]: A Meta-Analysis and Implications for Paleo-CO2 Reconstruction. Plant Ecophysiol 1 (2025) doi:10.53941/plantecophys.2025.100001.

45. Pillitteri, L. J., Sloan, D. B. & Torii, K. U. Termination of asymmetric cell division and differentiation of stomata. Nature 445, 501–505 (2007).

46. Miao, Z. et al. Easy Dataset: A Unified and Extensible Framework for Synthesizing LLM Fine-Tuning Data from Unstructured Documents. Preprint at 10.48550/arXiv.2507.04009 (2025).

47. Purcell, S. et al. PLINK: A Tool Set for Whole-Genome Association and Population-Based Linkage Analyses. The American Journal of Human Genetics 81, 559–575 (2007).

48. Yu, J. et al. A unified mixed-model method for association mapping that accounts for multiple levels of relatedness. Nat Genet 38, 203–208 (2006).

