## Supplemental Table 1-3 for "EpiReasoner: An Integrated Artificial Intelligence Framework for Phenotype-to-Genotype Reasoning in Plant Epidermal Development"

**Table S1. Summary of publicly available open-source stomatal and leaf epidermis image datasets used in this study.**

| Dataset name | Nation | Method | Amount of Raw Data | Source |
|---|---|---|---|---|
| Broad Bean & Wheat Stomata | China | Epidermal Peel | 160 + 951 | https://doi.org/10.5281/zenodo.6302925 |
| LeafNet | China | Epidermal Peel | 836 | https://leafnet.whu.edu.cn/suppdata |
| wheat_dataset | China | Epidermal Peel | 1000 | https://github.com/WeizhenLiuBioinform/stomatal_index |
| 278 Vascular Plant | UK | Epidermal Peel | 1556 | https://datadryad.org/dataset/doi:10.5061/dryad.g4q6pv3#usage |
| Populus & 17 Hardwood Species | USA | Impression | 10715 | https://doi.org/10.1038/s41597-023-02657-3 |
| Cultivated Sunflower | USA | Impression | 7288 | https://datadryad.org/dataset/doi:10.5061/dryad.xgxd254np |
| Mangrove Forest&Swamp Forest | Bangladesh | Impression | 1063 | https://data.mendeley.com/datasets/4brcwhmvyk/3 |
| Stoma_Saff | Italy | Impression | 138 | https://www.kaggle.com/datasets/claudiorusso9/stoma-saff |
| Maize | Brazil | Impression | 200 | https://zenodo.org/record/3938047#.YE_l6v7Q85k |
| Casado-Garcia2020_LabelStoma | Spain /USA | Impression | 1800 | https://github.com/ancasag/labelStoma?tab=readme-ov-file#datasets-and-weights |
| Cuticle database | USA | Fossil | 1374 | https://cuticledb.eesi.psu.edu/ |
| Ginkgo | USA | SEM | 608 | |
| Populus balsamifera | USA | DIC | 3298 | |
| USNM/USBG | USA | DIC | 1102 | |
| StomataScorer | China | In-Situ | 2500 | http://plantphenomics.hzau.edu. cn/download_checkiflogin_en.action |
| StomataSeg | Australia | In-Situ | 318 | StomataSeg: Semi-Supervised Instance Segmentation for Sorghum Stomatal Components |

**Table S2. Evaluation metrics for assessing instance segmentation performance of stomatal and pavement cells.**

| Metric Name | Symbol | Mathematical Definition | Parameter Description |
|---|---|---|---|
| Precision | $P$ | $\text{Precision} = \frac{\text{TP}}{\text{TP+FP}}$ | TP: True Positives (correctly predicted cell pixels);<br>FP: False Positives (background predicted as cell).<br>FN: False Negatives (cell pixels predicted as background). |
| Recall | $Rec$ | $\text{Recall} = \frac{\text{TP}}{\text{TP+FN}}$ | |
| F1-Score | $F1$ | $\text{F1} = \frac{2\times\text{Precision}\times\text{Recall}}{\text{Precision+Recall}}$ | |
| Dice Coefficient | $Dice$ | $\text{Dice} = \frac{2\times\text{TP}}{2\times\text{TP+FP+FN}}$ | |
| Mean Average Precision | $mAP$ | Multi-IoU threshold integration | Calculated via standard COCO API over multiple Intersection over Union (IoU) thresholds. |
| Mean Average Recall | $mAR$ | Multi-IoU threshold integrationI | Calculated via standard COCO API over multiple IoU thresholds. |
| Aggregated Jaccard Index | $AJI$ | $\text{AJI} = \frac{\sum_{i=1}^{N} \lvert G_i \cap P_j \rvert}{\sum_{i=1}^{N} \lvert G_i \cup P_j \rvert + \sum_{k\in U} \lvert P_k \rvert}$ | $G_i$: $i$-th ground truth instance; $P_j$: Predicted instance with maximum IoU to $G_i$; $U$: Set of unmatched predicted instances (false positives); $N$: Total number of ground truth instances. |
| Boundary IoU | $BoundIoU$ | $\text{BoundIoU} = \frac{\lvert B_{\text{G}}^{(d)} \cap B_{\text{P}}^{(d)} \rvert}{\lvert B_{\text{G}}^{(d)} \cup B_{\text{P}}^{(d)} \rvert}$ | $B_{\text{P}}$, $B_{\text{G}}$: Dilated ground truth and predicted boundary pixel sets within Euclidean distance threshold $d$. |
| Normalized Surface Distance | $NSD$ | $\text{NSD} = \frac{\lvert B_{\text{P}} \cap D_\tau(B_{\text{G}}) \rvert + \lvert B_{\text{G}} \cap D_\tau(B_{\text{P}}) \rvert}{\lvert B_{\text{P}} \rvert + \lvert B_{\text{G}} \rvert}$ | $B_P$, $B_G$: Boundary pixel sets; $D_\tau(\bullet)$: Distance transform function within tolerance $\tau$ (set to 2 pixels). |
| 95% Hausdorff Distance | $HD95$ | $\text{HD95} = \max(d_{95\%}(B_{\text{P}}, B_{\text{G}}), d_{95\%}(B_{\text{G}}, B_{\text{P}}))$ | $d_{95\%}(X, Y)$: The 95th percentile of the shortest Euclidean distances from all points in set $X$ to set $Y$. |

**Table S3. Phenotypic traits of stomatal complexes and pavement cells with their calculation methods.**

| Metric Name | Symbol | Calculation Method / Equation | Parameter Description |
|---|---|---|---|
| **I. Stomatal Complex Morphological Features** | | | |
| Stomatal Area/Perimeter | $A_{\text{stoma}}$ $P_{\text{stoma}}$ | Pixel integration & curve length | Both metrics are measured on the physically calibrated polygon contour of the entire stoma. |
| Feret's Diameter | $D_{\text{F}}$ | Maximum caliper distance | The longest distance between any two parallel tangents bounding the stomatal contour. |
| Major/Minor Axis Length | $L_{\text{major}}$ $L_{\text{minor}}$ | Eigenvector length calculation | Extracted via Principal Component Analysis (PCA) of the coordinate covariance matrix of the stomatal contour. |
| Spatial Deflection Angle | $\theta$ | $\theta = arctan\left(\frac{v_y}{v_x}\right)$ | Angle between the major axis directional vector and the horizontal axis of the reference coordinate system. |
| Pore Area | $A_{\text{pore}}$ | Pixel integration | The physical area of the internal opening region. |
| Pore Major/ Minor Axis Length | $L_{\text{pore\_major}}$/ $L_{\text{pore\_minor}}$ | Eigenvector length calculation | Extracted via PCA specifically applied to the internal pore region. |
| Aperture Index | $I_{\text{aperture}}$ | $I_{aperture} = \frac{L_{pore\_minor}}{L_{pore\_major}}$ | $L_{pore\_minor}$: Pore minor axis; $L_{pore\_major}$: Pore major axis. Represents stomatal opening status. |
| Pore Area Ratio | $R_{\text{pore}}$ | $R_{pore} = \frac{A_{pore}}{A_{stoma}}$ | $A_{pore}$: Internal pore area; $A_{stoma}$: Total stomatal complex area. |
| **II. Pavement Cell Morphological Features** | | | |
| Pavement cell Area/Perimeter | $A_{\text{PC}}$ $P_{\text{PC}}$ | Pixel integration & curve length | Both metrics are measured on the physically calibrated polygon contour of the entire pavement cell. |

| | | | |
|---|---|---|---|
| Circularity | $C$ | $C=\frac{4\pi A}{P^2}$ | $A$: Cell pixel area; $P$: Cell contour perimeter. |
| Solidity | $S$ | $S=\frac{A}{A_{hull}}$ | $A$: Cell pixel area; $A_{\mathrm{hull}}$: Area of the cell's minimum convex hull. Quantifies lobe indentation. |
| Rectangularity | $R$ | $R=\frac{A}{A_{MBR}}$ | $A$: Cell pixel area; $A_{MBR}$: Area of the Minimum Bounding Rectangle. |
| Neck Width | $W_{\mathrm{neck}}$ | $W_{neck}= min\,\|\|p_{defect\_i}-p_{defect\_j}\|\|_2$ | $p_{defect\_i}$, $p_{defect\_j}$: Spatial coordinates of two opposite convexity defect points (lobe indentations). |
| Lobe Length | $L_{\mathrm{lobe}}$ | $L_{lobe}=\|\|p_{tip}-p_{midpoint}\|\|_2$ | $p_{tip}$: Lobe tip (local curvature maximum); $p_{midpoint}$: Center point of the corresponding neck baseline. |
| Protrusion Depth | $D_{\mathrm{protrusion}}$ | Orthogonal projection distance from the lobe tip to the neck baseline (line connecting two adjacent defect points) | Orthogonal distance from $p_{tip}$ to the line connecting two adjacent convexity defect points. |
| **III. Spatial Topology and Distribution Features** | | | |
| Stomatal Density | SD | $\mathrm{SD}=\frac{\mathrm{Count(stoma)}}{A_{\mathrm{tissue}}}$ | Count(stoma): Total number of stomata in the analyzed region. $A_{\mathrm{tissue}}$: Total physical area of the analyzed epidermal tissue region. |
| Stomatal Index | SI | $\mathrm{SI}=\frac{\mathrm{Count(stoma)}}{\mathrm{Count(stoma)+Count(PC)}}\times 100\%$ | Count(stoma): Total number of stomata; Count(PC): Total number of pavement cells in the same region. |
| Shortest Topological Path | $D_{\mathrm{topo}}(s_i, s_j)$ | $D_{\mathrm{topo}}(s_i, s_j) = \min_{\mathrm{path}}\{\mathrm{Count(pavement\_cells)} \in \mathrm{path}(s_i, s_j)\}$ | $s_i$,$s_j$: Nodes representing adjacent stomata; Count(PC): Number of pavement cells traversed in the Breadth-First Search (BFS) graph. |
| Stomatal Cluster Violation Rate | $SCVR$ | $\mathrm{SCVR} = \frac{\|\{(s_i, s_j) \in E \| D_{\mathrm{topo}}(s_i, s_j) = 0\}\|}{\|E\|}\times 100\%$ | $E$: Total set of stomatal adjacency edges defined by Delaunay triangulation; $D_{\mathrm{topo}} = 0$: Condition indicating a direct physical contact violation. |
| Tricellular Junction Density | $\rho_{\mathrm{TCJ}}$ | $\rho_{\mathrm{TCJ}} = \frac{\mathrm{Count(TCJ)}}{A_{\mathrm{tissue}}}$ | Count(TCJ): Total number of tricellular junctions. $A_{\mathrm{tissue}}$: Total area of the analyzed epidermal tissue region. |
